# Calculating likelihoods and likelihood ratios at SNPs-based mixtures. A reappraisal of the binomial inference, as applied to forensic identity tests

**DOI:** 10.1101/2021.02.08.430218

**Authors:** Vincenzo L. Pascali

## Abstract

Single nucleotide polymorphisms (SNPs) are useful forensic markers. When a SNPs-based forensic protocol targets a body fluid stain, it returns elementary evidence regardless of the number of individuals that might have contributed to the stain deposition. Therefore, drawing inference from a mixed stain with SNPs is different than drawing it while using multinomial polymorphisms. We here revisit this subject, with a view to contribute to a fresher insight into it. First, we manage to model conditional semi-continuous likelihoods in terms of matrices of genotype permutations vs number of contributors (NTZsc). Secondly, we redefine some algebraic formulas to approach the semi-continuous calculation. To address allelic dropouts, we introduce a peak height ratio index (‘h’, or: the minor read divided by the major read at any NGS-based typing result) into the semi-continuous formulas, for they to act as an acceptable proxy of the ‘split drop’ (Haned et al, 2012) model of calculation. Secondly, we introduce a new, empirical method to deduct the *expected* quantitative ratio at which the contributors of a mixture have originally mixed and the *observed* ratio generated by each genotype combination at each locus. Compliance between observed and expected quantity ratios is measured in terms of (1-χ^2^) values at each state of a locus deconvolution. These probability values are multiplied, along with the h index, to the relevant population probabilities to weigh the overall plausibility of each combination according to the quantitative perspective. We compare calculation performances of our empirical procedure (NITZq) with those of the EUROFORMIX software ver. **3.0.3**. NITZq generates LR values a few orders of magnitude lower than EUROFORMIX when true contributors are used as POIs, but much lower LR values when false contributors are used as POIs. NITZ calculation routines may be useful, especially in combination with mass genomics typing protocols.

## Introduction

In forensic genetics, a long-standing tradition of good practice has endorsed the adoption of multi-allelic polymorphisms (essentially, short tandem repeats; STRs) for the routine casework. The assumption underlying this choice is that a handful of these markers can assemble effective protocols and help to identify the donor(s) of a stain left at the crime scene. However, single nucleotide polymorphisms (SNPs) are recently qualifying as an prominent class of polymorphisms to adopt in the forensic work.

SNPs are mostly bi-allelic, and the Mendelian distribution of their genotypes (the algebraic expansion of a pair of gene frequencies) is easy to determine by just sampling a small population of unrelated individuals. SNPs are also ubiquitous, and a typical random human genome differs from the reference human genome by 4.1 - 5.0 million sites most of which are SNPs. [1]. *Last not least*, the next generation sequencing procedures (NGS) today used to type SNPs are not affected by most of those technical artefacts that normally complicate the process of STR typing. In view of this favourable background [2], several SNPs panels have been assembled for human identification and validated for the forensic routine [3–20].

However, the everyday application of these SNPs panels to cases of criminal identification has proceeded at slower pace than expected, possibly due to a series of circumstances:

a. SNPs protocols enlist more numerous markers compared to STRs; readapting calculation procedures / software routines (either based on the qualitative /semi-continuous approach [21] or on the quantitative approach [22–27]) to hundreds -thousands loci has proved to be more laborious and time consuming than initially expected;
b. Stains of forensic interest are mostly made of an admixture of multiple contributors; when a SNPs protocol is employed to type these mixed stains, the evidence tends to be uniformly biallelic and, based on this evidence, one cannot formally exclude any specific suspect. This arises the serious problem of discriminating falsely compatible results from genuine matches by a statistic approach and not by the formal outcome of the test.

As a consequence of this deadlock, and with some notable exception [28], the routine of criminal DNA tests still draws on without SNPs in several Countries.

The computational bottleneck at SNPs mixtures analysis has recently become the object of several studies. Estimates based on absolute likelihoods have been tried and proposed as the computational complement to mass typing protocols (Homer et al (2008) [29]; Voskoboinik and Darvasi (2011) [30]), but this ‘frequentist’ approach (further developed by Isaacson et al (2015) [31]) has arisen criticisms (Clayton, 2010; [32]; Egeland et al 2012[33]). Furthermore, Voskoboinik et al 2015[34] have shown how a single contributor can be identified in mixtures by calculating true likelihood ratios at a protocol based on 3,000 SNPs (0.002<P(MAF) < 0,25. Nearly at the same time, Gill et al (2015) [35] have explored the feasibility of the semi-continuous (or qualitative) calculations to the case of 2-person and 3-person mixtures. More recently, Bleka et al [36] have readapted the open-source *EUROFORMIX* software to the biallelic scheme and performed quantitative calculations starting from the ‘reads’ produced by a massive parallel sequencing (MPS) equipment.

In this paper we will revisit the logic of calculating binomial likelihoods and LRs at forensic mixtures, under both the semi-continuous and the quantitative perspective.

## Materials and methods

### Semi-continuous calculations

#### Matrices to calculate likelihoods as at SNPs mixtures

At any mixture, the trace evidence (Et) contains n contributors (n>1; this is the whole lot of individuals involved in a crime scene *and* leaving cells into the biological trace in matter). With binomial SNPs, the evidence may only be single-allele (Et{a} or double-allele Et{a,b}. When the number of contributors increases, the double-allele evidence prevails and the mono-allelic becomes an exception to find. If n is the only prior assumption to formulate on the mixture content, Et{a,b} generates a matrix with as many columns as the number of contributors and as many rows as the number of genotype permutations that can as equally express that evidence - whereas (Et{a} remains with no matrix at all. Such matrices have been denominated ‘deconvolutions’. Deconvolutions of multinomial STR loci have been explored by Lynch and Cotton (2018) [37] and these authors have warned on the numerosity and complexity of STRs deconvolutions generated by dozens of genotypes and more than three contributors. This warning, however, does not apply to SNPs: as opposed to STRs, SNPs have just one Et{a,b} deconvolution per contributor, no matter how numerous the contributors are. Moreover, the number of records at each of these binomial deconvolutions remains acceptably low even the number of contributors is high. In fact one can count 7 permutation rows (or deconvolution states) at the 2PM case, 25 rows at the 3PM deconvolution, 79 at the 4PM, 249 at the 5PM, 727 at 6 PM, 2185 at 7PM and 6559 at 8PM. The two circumstances (one matrix per class of mixture; a limited number of coexistent combinatorial records per matrix) make it convenient to pre-compile a handful of universal matrices (one per number of contributors), place them into as many Excel worksheets and use the worksheets for computing over all binomial loci of interest. Matrices may be made universal by adopting always the same allele nomenclature - A for the quantitatively predominant allele, B for the other; AA, AB and BB for the relevant genotypes - for whatever locus. If used for semi-continuous calculations, matrices will only contain genotype frequencies, so that using the universal matrix for the n^th^ SNP is only a matter of updating the worksheet’s pair of gene frequencies with those of the locus of interest. Universal matrices become convenient to use when the number of SNPs placed at a forensic protocol is counted in hundreds [17–20] or possibly thousands [16, 30, 34]).

When further prior hypotheses are formulated introducing η persons of interest (POIs; known contributors, or Ks; 1 ≤ η ≤n) the inference to draw becomes conditional to both n and η (Benshop et al 2015; [38]). Just as noticed for Et, sets of POIs can have a single-allele pattern Kt{a} or a biallelic pattern Kt{a,b}. POIs sets are used as a condition to single out subsets of the general matrix containing their genotype (see Appendix). This may be achieved by embedding a few, simple *‘if-then’* excel conditional formulas into the relevant Excel worksheet. These subsets are themselves true likelihoods and their arithmetical value is inferred by summing the permutation states products contained at only those records that contain the known genotype. We here supply seven precompiled spreadsheets (to be downloaded from https://www.vincepascali.it/docs/NITZsc_2_to_8_contrs.xlsx), one per number of contributors (from 2PM to 8PM), each allowing computations that allocate up to two known genotype per matrix.

#### Calculating likelihoods by algebra

When compared one to another, the evidence and the POI allelic subsets generate nine assortments, shown in Fig 1. Fig 1 contains matching cases and exclusions. Exclusions only emerge when the evidence is made of one-allele. When this happens, either K and Et are disjointed sets (Kt ⋂ Et=Ø; no allele in common; for example Et{a} comparing to Kt{b}) and viceversa) or Et is a subset of Kt: Et ⊂ Kt (whereas in the case of compatibility one would normally expect the reverse Kt ⊂ Et to be true).

**Fig 1.**
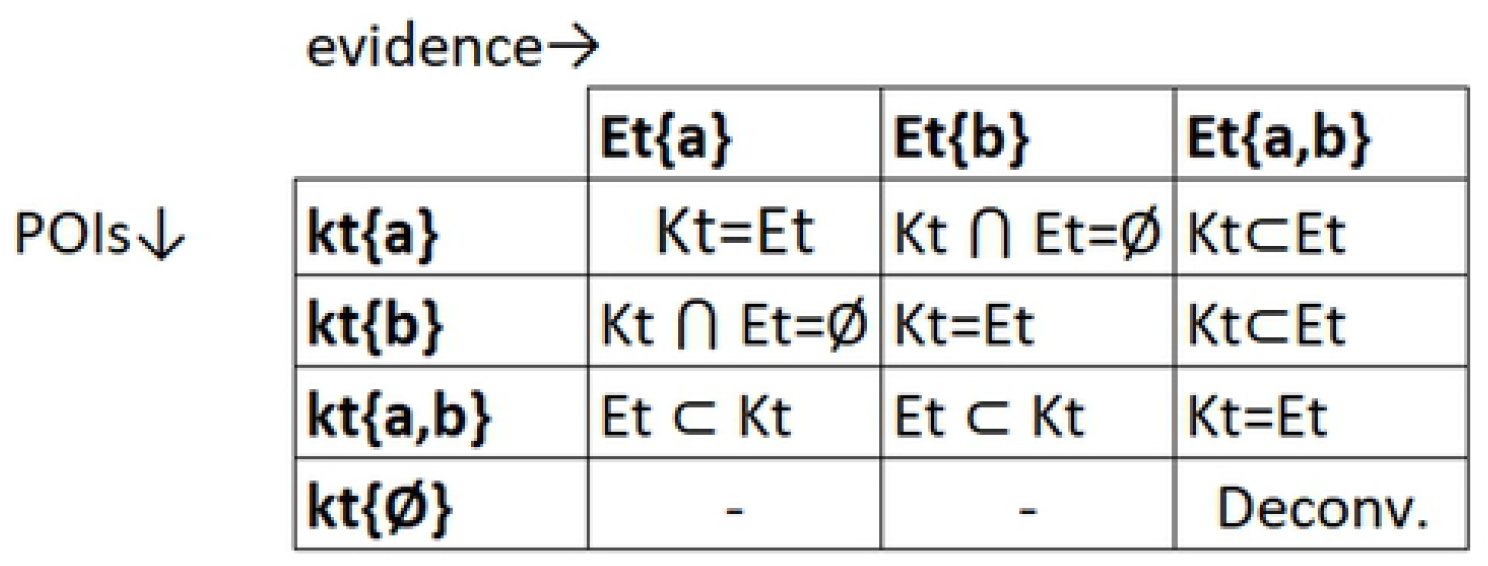
Assortments of the evidence and the POI allelic subsets. This matrix shows that there is a limited number of outcomes (just nine, in fact) to expect at a SNPs based identity test.

All of these nine cases may be worked out by algebra. Principles ruling the algebra are:

1. at a single-source trace, the whole evidence is: PEt (a+b)^2^ = P(E_t_{a})+ P(E_t_{b})+ P(E_t_{a,b})=1;
2. the generalization of the single-source trace model to n contributors requires exponentiation to n of all members of the evidence: P (a+b)^2n^ = P(a^2n^)+ P(b^2n^) + P(a-b)^2n^ [21];
3. introducing η obliged contributors and calculating relevant likelihoods involves to subtract η to the original exponent: P(a+b)^2n^ → P(a+b)^2(n-η)^.

#### Algebraic notations for compatible semi-continuous likelihoods

The following conditional formulas apply to compatible cases of table 1:

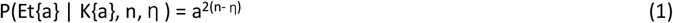

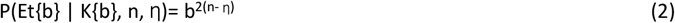

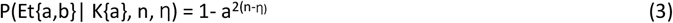

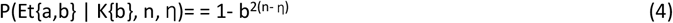

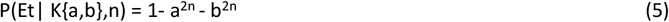

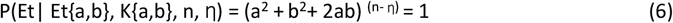

where a,b in the right side of the equations are the gene frequencies of the homonymous SNPs alleles. These formulas are a generalization of those reported by Voskoboinik et al 2015 [34] – modified by introduction of rule 3 of previous chapter. As previously noticed by these authors, the six formulas do not account for cases where there is a mismatch between the evidence and the POI.

#### Algebraic notations for incompatible likelihoods

We here assume that the mono-allelic evidence derives from an originally bi-allelic evidence by the loss of one allele (allele dropout). Within our nomenclature system, only the minor quantity allele can drop out of an evidence, with some probability (P(D)) that proportionally increases as soon as the template concentration decreases down to critical values. In the available literature P(D) values have been inferred from experiments conducted on various templates dilution, by logistic regression analysis (Tvedebrink et al 2009 [39]; and Buckleton et al 2014 [40]) – and these experimental values have been incorporated into semi-continuous calculations. A current computational approach relying on such dropout probabilities is the ‘split drop approach’, developed by Haned et al in 2012 [41] and readapted to the case of SNPs by Voskoboinik et al [[34]. The split-drop approach is actually adopted by the two widespread software LRMIX and EUROFORMIX [42].

We instead take the approach of incorporating dropout probabilities into a few algebraic formulas working out the task that is normally committed to the split drop approach:

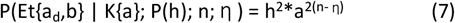

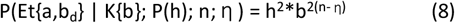

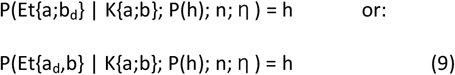

It is here to notice that, with reference to the notations adopted in figure 1, Et{a;b_d_} replaces the original Et{a} notation and Et{a_d_;b} replaces Et{b}, respectively. P(h) is a proxy for the experimental logit values. For its meaning, read on the following paragraphs.

#### Algebra for ‘dropped out and still compatible’ cases

Although the concept of dropout is invoked to account for cases of contributors that do not match the evidence, ‘exclusion’ and ‘dropout’ are not logically equivalents. In fact there may be instances where an allele drops out of the evidence and the resulting reference remains compatible to the POI. Such ‘latent’ dropout cases can be modelled by turning equations 1 and equ. 2 into:

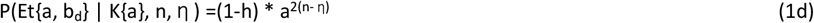

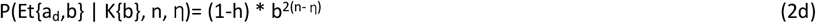

It is to note that, if using our conventional nomenclature (a, major read allele; b, minor read allele) equation 2d is never actually used.

#### Complete algebraic inference

The algebraic solutions we have just introduced (1d; 2d; 3-6; 7-9) can now cover all the inferential patterns expected to show up at any test based on binomial markers. An overview of the nine relevant cases is in Fig 2.

**Fig 2.**
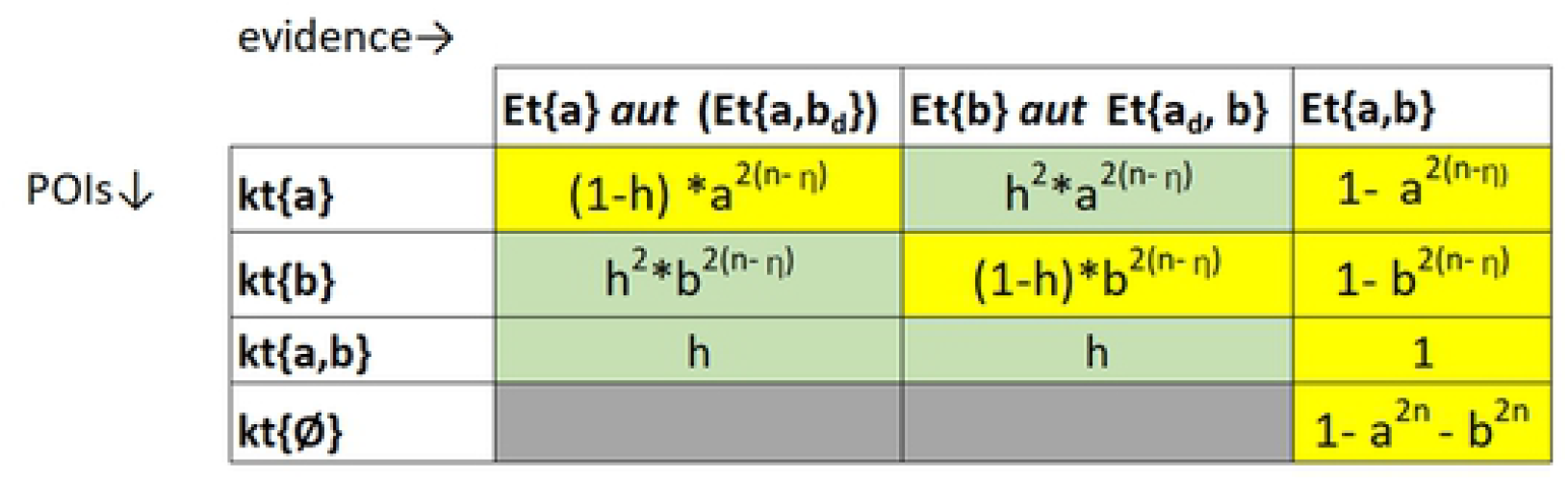
Algebraic solutions given to all possible evidence/POI allelic subsets assortments of Fig1. The mono-allelic evidence appearing in Fig 1 is reinterpreted under the assumption that dropout events may have altered Et{a, b}. Compatible cases are highlighted in yellow background, exclusions in green background.

#### The h index as a proxy for experimental P(D) values

Dropout occurs at SNPs too [43], but specific logit dropout values have not been so far exploited for SNPs. This is not surprising: these values should be inferred (by experimental data logit treatment) from *each locus in use at a given protocol- possibly by each laboratory adopting the protocol*. And because forensic protocols are often made of hundreds / thousands loci, this goal is not easy to attain in practice. That being the case, a good deal of semi-continuous calculations at SNPs mixtures (as those performed by the typical users of LRMIX and EUROFOMIX) end up to rely on arbitrary P(D) values. This encourages to search for other solutions.

We found the a convenient proxy for the P(D) is the h index, introduced by Kelly et al [44]), in its easiest form (the minor red simply divided by the major read). There are several reasons to introduce this index in place of logit-derived P(D) values:

a. h applies as well to single-source stains and mixtures;
b. at mixtures, h may represent the whole propensity of the minor-read allele to drop out of the mixed evidence;
c. (1-h) can estimate the probability of existence of the two-allele evidence;
d. The equivalence h ≈ P(D) has a theoretical justification: it has been predicted that a P(D) can automatically emerge from the peak height analysis, without any external logistic regression modelling (Cowell, 2018 [45]); if starting from this standpoint, a typical P(D) value to deduct from the peak area analysis cannot be but the h index.

Index h enters into the likelihood formulas as shown in the previous paragraphs.

#### Dropins

According to an official definition issued by the ISFS commission [46], dropin is an allele that is not associated with the crime sample and remains unaccounted both under the prosecutor and the defence hypothesis. In principle, a dropin phenomenon is nothing more than an additional, ‘uninteresting’ contributor to a mixture and its case could be modelled by increasing the number of contributors. Since the SNPs-based evidence generates relatively simple deconvolutions even at high numbers of contributors we address the dropin case by systematically switching the current scheme of computation to one scheme with one more contributor. We will later see that this has limited effect on the LR value.

#### Number of contributors

Guessing on the number of contributors that enter in a SNPs mixture may be difficult. The importance of the whole issue is directly proportional to the impact the value of n may have over the LR values. Bleka et al 2017 [36] have explored this issue by simulating 100 mixtures with two to up to six virtual contributors and calculated likelihoods at various quantitative POI proportions to the POI. They found that, when the POI is not quantitatively predominant, their two-person mixture model of analysis would return them higher LRs compared to the three- or more-person models. They also found that, from three-persons mixtures onwards, increasing the number of contributors has a negligible effects over the POI’s LR, regardless of the POI’s quantitative predominance. We have an explanation to this phenomenon: unlike higher-rank mixtures, 2-persons mixtures cannot have permutation states with full assortment of binomial genotypes (AA+AB+BB). Switching from 2PM to 3PM has therefore the effect of completing the genotype assortment, elongating the deconvolution, increasing the number of states and lowering the MR. The same does not occur while switching from 3PM to 4PM or to higher-rank models. There is consequently much logic in believing that 3PM is the minimum standard to endorse at binomial mixture analysis. In this paper, however, we will always calculate LRs under three distinct hypotheses (2PM, 3PM, 4PM).

#### LRs

Each likelihood in figure 2 may represent the viewpoint of the prosecution or the defense at a typical judicial test. A ratio between two of these likelihoods will return the value of the available, prevailing evidence. The two parties may choose to opt for the same n or different n values but they usually diverge on the choice of the η value. The prosecutor’s choice η1, placed at the numerator of a ratio, is always higher than the defenses’ choice η2 - placed at the denominator ((η1- η2) ≥ 1). For what matters in this paper, the classical LR formula [47] is therefore written as:

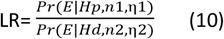

### Quantitative calculations

#### Background

Algebra has no power on quantitative calculations because all states of a quantitative deconvolution have unique probability values and algebraic simplifications cannot operate on these matrices. Leaving out algebra and staying with matrices is therefore our way to tackle the quantitative issue. (The central role of matrices in addressing quantitative mixture analysis is in our view unsurprising, as this kind of analysis has been extensively addressed by adopting Bayesian networks, which are a tool enabling to harness complex matrices with comparatively little effort [48, 49]).

The issue of quantitative mixture analysis has been reviewed several times [50–54] and – according to most reviewers - a key issue in the field is finding a suitable way to infer the peak fraction values pertaining to each contributor from the whole of the mixture evidence. In a pioneer-paper, Cowell et al [55] used the gamma distribution to deduct all individual peak heights fractions that coalesce to shape the mixed STRs profile. Almost at the same time, Curran (2008; [23]) introduced Markov Chain Monte Carlo simulations to infer this distribution. These two methods have earned much momentum in the subsequent decade and today many dedicated software pieces are based on the gamma model or MCMC methods. Here, however, we will adopt a new, all empirical method for deducting the mixing ratio at a mixture.

#### An empirical method enabling to capture MR fractions

Deducting the ratio at which contributors mix a mixture is difficult because contributors tend to share alleles and to overlap within the evidence. This phenomenon is heavier when numerous contributors concur to produce the same trace and at binomial loci.

However, at certain states of a mixture deconvolution, a single contributor is able to give the whole of the reading signal pertaining to a single allele. At each of these non-overlapping genotypes (NOGs), the number of reads referring to the NOG, divided by the sum of all reads, returns the proportion at which the NOG-bearing contributor has entered in the mixture.

At SNPs there also exist some simple-overlapping genotype combinations (SOGs) enabling to deduct the individual contribution with additional but elementary computation. Binomial NOGs and SOGs combinations are shown in Fig 3, along with the corresponding formulas to adopt for extracting the mixing ratio. When no POI is available, NOGs and SOGS are of no use. In this case, all genotypes will permute within each position in the matrix, each genotype will end up to belong to each contributor and all contributors will share the same quantity (0,5 to 0,5 at the 2PM; 0,33: 0,3 : 0,33 at 3PMs; 0,25×4 at 4PMs etcetera). But when a NOG/ SOG bearing POI becomes available, NOGs/SOGs will no longer permute and the reconstruction of the POI quantity becomes possible. Unknown contributors will of course continue to permute and share identical fractiond of the remaining quantity. To make a numerical example: at a 3PM a POI amounting to 0,4 /1 will leave a 0,6 /1 to the two unknowns who will amount to 0,3 /1 each. When the POI is AA, NOGs and SOGs values generated from all AA (BB)n and AA (AB)n series will correspond to the same AA POI and they become equally possible values. Both values ar then considered for deriving the MR. A POI may be a true contributor or a false contributor. If true, his/her quantitative NOGs will cluster around just a definite value, net of measurements errors. If he is a false contributor, its quantitative measurements will flatten on the 1/n value. In turn, MRs extracted from true contributors will return high LR values, false contributors will have LR ≤1. When true contributors have an MR proportion close to the 1/n value, they behave as if they were false contributors ( LR falls below 1). If more the one POI is available, the respective NOGs can be extracted either separately or together. Because SNPs protocols are usually made of some hundreds loci, a significant number of NOGs and SOGs are usually intercepted at any given experiment.

**Fig 3.**
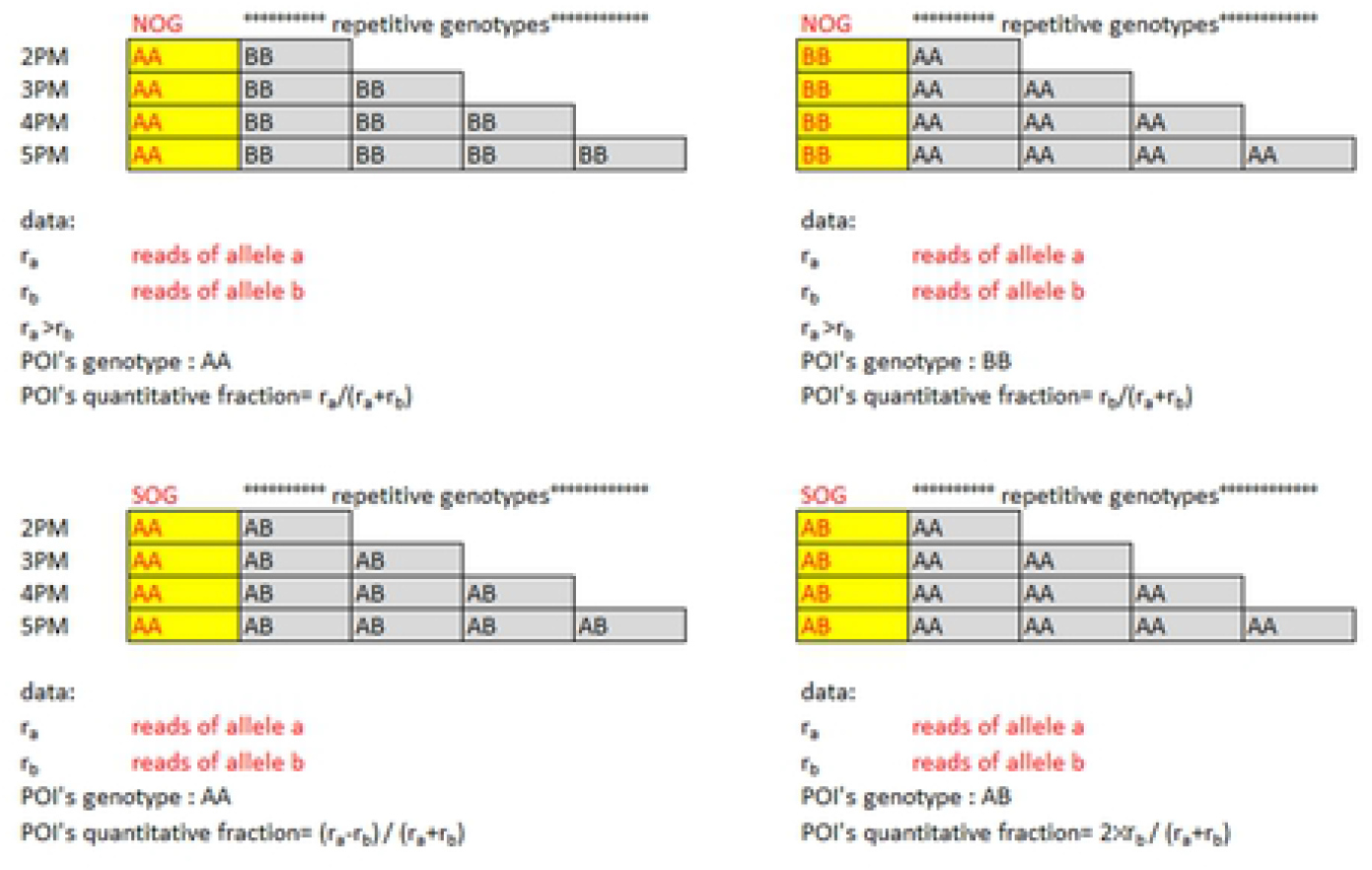
Non overlapping (NOCs) and simple overlapping (SOCs) combinations at SNPs. Genotypes to adopt for calculating the POI’s mixing ratio are in red typeface; repetitive, non useful genotypes in black. Increasing the number of contributors leads to an elongation of the non-useful genotypes whereas NOGs and SOGs kept outside the overlapping zone and remain unique. Data and the formulas to adopt for calculating the MR are specified under the relevant scheme.

Proportions deducted from NOGs and SOGs can be generalized to all loci (either overlapping or non-overlapping) and become a proxy value for the general mix ratio (MR).

#### Introducing empirical MR values into probability calculations

Once deducted from the NOGs/SOGs statistics, the empirical MR become the ‘espected’ value to retrieve at every single combination of genotypes entering in a likelihood matrix. ‘Observed ‘ values of mixing ratios are in turn deducted from each, n^th^ deconvolution state of the mixture, by simple algebraic operations reproducing those in figure 3 and placed within our worksheets (to be downloaded at: www.vincepascali.it/docs/NITZq_SNPs_2PM.xlsx; www.vincepascali.it/docs/NITZq_SNPs_3PM.xlsx; www.vincepascali.it/docs/NITZq_SNPs_4PM.xlsx).

Observed and expected MR values are compared by setting series of chi square tests. The (1-χ^2^) value deducted at each individual state will eventually return the probability of that state to exist in the mixture.

#### Three indexes for each likelihood value calculation

Our three NITZq matrices distribute three statistical indexes: the (1-χ^2^) MR values, the h index (the minor read divided by the major read) and the population genotype frequency. These three values embody as many independent properties of mixtures (the MR belongs to the specific mixture; the h index is an autosomal property; the genotype frequency is from the population) and they multiply at the level of each deconvolution state. All products are in turn summed to give the value of any likelihood of interest.

#### Flow chart of NITZ calculations

Although our NITZq worksheets have no pretence to make an integrated software, each flowcharts a whole series of 130-some SNPs Mendelian and quantitative data into a unique, semi-automatic procedure that return perfectly reliable LR calculations.

The sequence of passages included within NITZq worksheet flow chart is as below.

I. **evidence input**. The appropriate worksheet of the NITZ library (NITZq 2PM, NITZq 3PM and NITZq 4PM) is to be selected. Each worksheet has an *“input & MR”* spreadsheet that allows the user to input the following data (from the worksheet’s top-left side onwards and in sequential order per each locus): locus name; major read; minor read; gene frequency of the major read gene; gene frequency of minor read gene;
II. **POI genotypes input**; all POI(s) genotypes have first to be code-classified as AA (homozygous major read), BB (homozygous minor read), AB (major-minor heterozygous) according to the conventional, major-read allele / minor-read allele code. Code-classified genotypes are then paste into the input area as ‘reference data’ placed adjacent to the ‘evidence data’. This will trigger hundred calculations into a series of cross-referenced worksheets.
III. **basic calculations**: specific algebraic functions placed in a *“calculations”* worksheet calculate genotype frequencies, individual quantities and heterozygous ratio (h) at every record corresponding to each state of the loci deconvolutions (presently NITZq allocates 133 loci).
IV. **calculation of the MR**: at the “input & MR” spreadsheet, NOGS/SOGs values are intercepted based on the POI genotypes. Their values are summed and averaged and the quantity fraction belonging to the POI is deducted. Finally, (1-Q_POI_) divided by the number of unknowns will give the other fractions of MR. The array of the MR values is cross-referenced into thousands positions at the *‘calculations’* worksheet.
V. **(1- χ^2^) calculation:** the individual quantities deducted at each deconvolution state are compared to the NOGs/SOGs MR and transformed into (1- χ^2^) values (one value per record);
VI. **likelihood calculations**: genotype frequencies, h values and (1- χ^2^) values are multiplied across each deconvolution state and their products summed together to give the likelihood of choice
VII. **calculation of the LR** Likelihoods and LR calculations update instantly as soon as the evidence and POI input is pasted into the “input & MR” spreadsheet. Computation time is virtually zero.
VIII. **correction for incompatible cases** Extremely imbalanced read pairs in the evidence may indicate that all the underlying mixture contributors are homozygous and the evidence itself may qualify as single-allele (Et{a}). Because no deconvolution and no quantity calculation applies to all tests bearing the Et{a} evidence, these cases need to be identified and brought back to algebra. To spot cases as these, we calculate the h index at every locus evidence, set a threshold for the h-evidence value below which the locus is declared ‘single-allele’ and reclassify. Spotting single-allele evidence is essential in order to identify privileged Evidence/Reference assortments: Et{a} vs POI_AA_ Et{a} vs POI_BB_ Et{a} vs POI_AB_.

Generally, the first assortment tends to give high LRs placed well above 1, the latter two low/very low LRs.

We indiscriminately apply this correction to the semi-continuous and the quantitative scheme.

### Testing datasets

To test the performance of our formulas, matrices and worksheets and compare it with high quality software we chose to download the “dataset 4: SNPs data” made available by Dr Ø. Bleka at the open-source EUROFORMIX website (http://www.EUROFORMIX.com/?q=data). This dataset contains evidentiary genotypic and quantitative data pertaining to several experimental 2PM mixtures and 3PM mixtures designed to cover various mixing ratios under accurately controlled conditions. ‘EUROFORMIX SNPs Data’ supplies typing results pertaining to a subset of 140 biallelic SNPs [56]. It also includes three POI genotypic dataset corresponding to three contributors (P1,P2,P3) that have been actually used to build the evidence. An array of European gene frequencies is also supplied. We singled out the XEN13_a (a two person mixture made of individual P1 and P2, mixing at five different MRs; 1:1; 1:3; 1:9; 3:1; 9:1) and the XEN46_a (a three person mixture made with individual P1, P2 and P3, mixing at: 1:1:1; 5:1.1; 1:5:1; 1:1.5; 5:5:1; 5:1:5; 1:5:5) as datasets for our comparison test. After elimination of incomplete typing data, a selection of 133 XEN13_a /XEN46 loci was eventually selected and brought to the statistical analysis. In addition, we used data from forty random Swedish individuals (the much valued courtesy of professor Andreas Tillmar, Department of Forensic Genetics and Forensic Toxicology, National Board of Forensic Medicine, Linköping, Sweden) as non-contributor datasets.

Each group of data drawn from this collection was thoroughly analysed by adopting the 2-person, 3-person and 4 person mixing scheme irrespective of the fact that the original evidence would originate from a two-person or a three person’s. All data were independently processed by using the EUROFORMIX shareware (version 3.0.3) and then by the collection of all NITZq worksheets.

## Results

### MRs from XEN 13a / XEN 46a, three true contributors as POI and a variable amount of unknowns

Fig 4 and Fig 5 show twelve series of mix ratio predictions as they come out from our NITZq 2PM, 3PM and 4 PM worksheets and from EUROFORMIX. The evidentiary datasets were an experimental three person mixture (XEN46a) containing three contributors P1,P3,P2 mixing at seven different ratios (1:1:1; 5:5:1; 5:1:5; 1:5:5; 5:1:1; 1:1:5; 1:5:1; quoted in the order P1,P3;P2) and a two-person mixture (Xen13a) containing two contributors P1 and P2 blending at five different balancing ratios (1:1;1;13;1:9; 3:1; 9:1; first quoted quantity is P1). P1, P3 and P2 were in turn individually used as person of interest along with one (KU), two (KUU) and three (KUUU) additional unknown contributors (KU; KUU; KUUU). The POI is here always a ‘true’ known contributor– i.e. a person whose DNA had been actually used to cast the mixture evidence - its genotypic array learned from the EUROFORMIX website. Just one of the three hypotheses on the number of contributors used to interrogate each evidentiary example was ‘true’ based on what’s declared in the EUROFORMIX website. The other predictions were assuming a wrong number of contributors. These data shows that NITZq quantitative POI predictions compare well to those issued by the EUROFOMIX software whereas predictions on unknown contributors operated by the two methods may occasionally diverge: in principle, NITZ always flattens the unknown MR proportions to exactly the same value; EUROFORMIX occasionally assigns differential values, other times distributes just one value to all unknowns. Predicting quantities under a wrong assumption about the true number of contributors has a limited effect on the effect on the POI’s share. In principle, expanding the contributors’ number shrinks the quantitative proportion left to a true POI.

**Fig 4.**
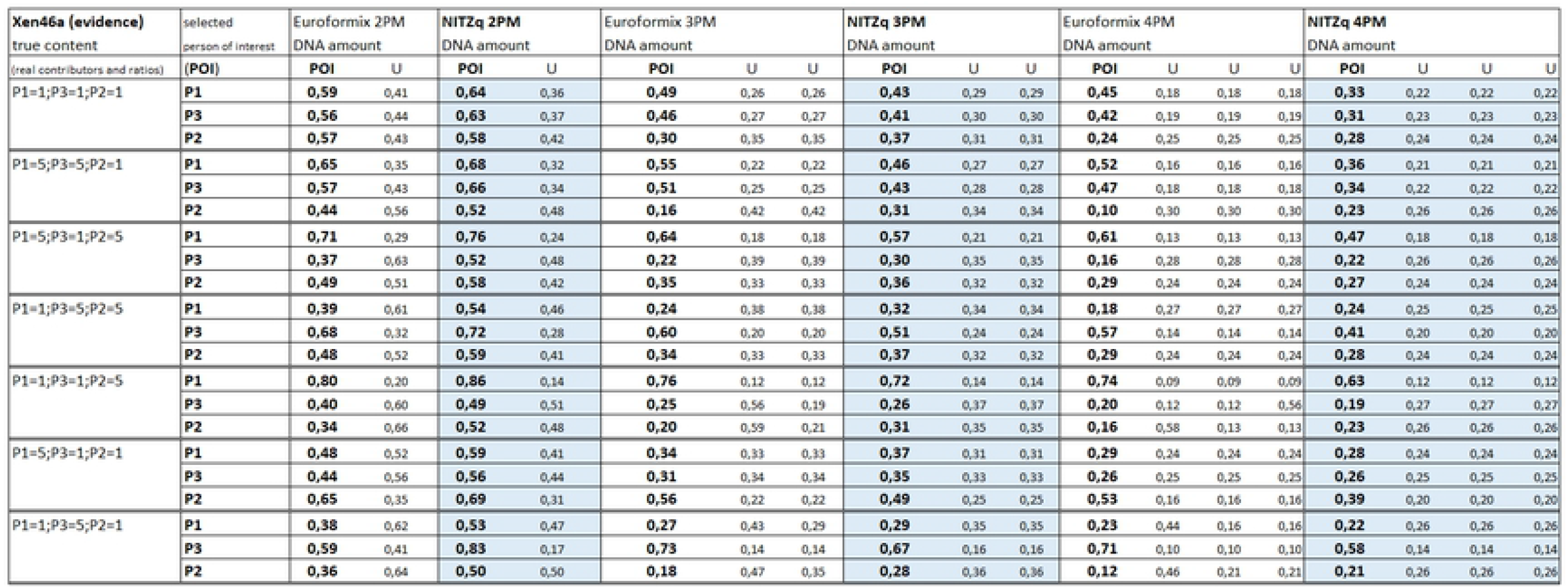
Amount of DNA assigned to a true contributor POI at a three-person-based experimental evidence (XEN46a)

**Fig. 5.**
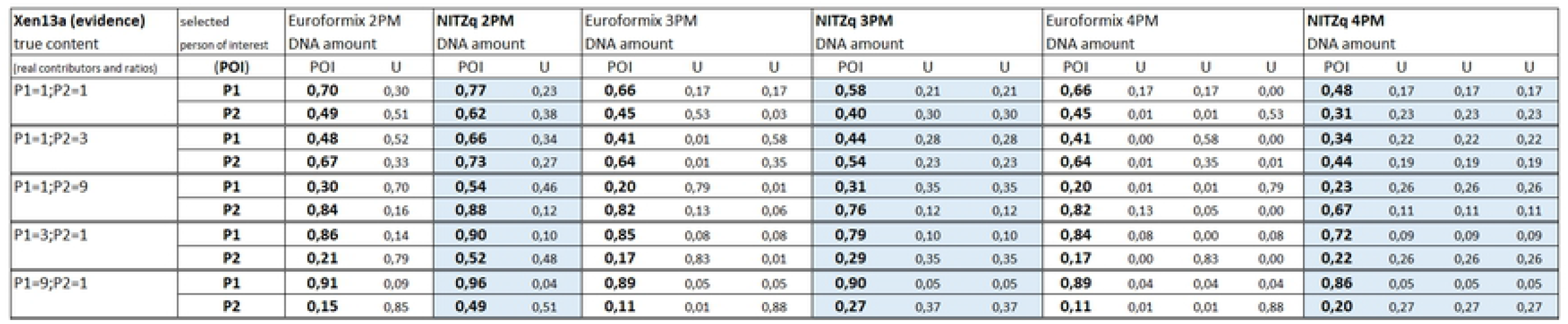
Amount of DNA assigned to a true contributor POI at a two-person-based experimental evidence (XEN13a)

### MRs, with false contributors in the role of POI

Another series of MR predictions were produced by replacing the ‘true’ P1,P3, P2 with false contributors in the role of POI. False contributors were taken one by one among a population of forty unrelated Swedes (A. Tillman’s data). Predictions followed the same scheme as before but for simplicity each POI was matched to a fewer evidentiary datasets (XEN13, ratios: 1:1; 1:9; XEN46a ratios: 1:1:1; 1:5:5; 1:1:5). With both NITZ and EUROFORMIX, quantitative fractions predicted to each of the forty POIs were so close one to another that they could be conveniently averaged with little standard deviation, as shown in Fig 6 and Fig 7. Figures here reflect the different approach of the two methods: EUROFORMIX seeks for the best fitting quantitative proportions in hundred random simulations, NITZ singles out values tending to 1/n, which is the result of a free deconvolution. We envision the 1/n value is eventually reached by predicting MRs over an infinite number of loci. Increasing the number of contributors has the effect of halving the quantitative fraction assigned by EUROFORMIX. NITZq moves in pretty much the same direction, but the decrease in the POI quantitative proportion has its floor in 1/n. Matching the same POIs to balanced or unbalanced mixtures has little effect on the predicted proportions.

**Fig 6.**
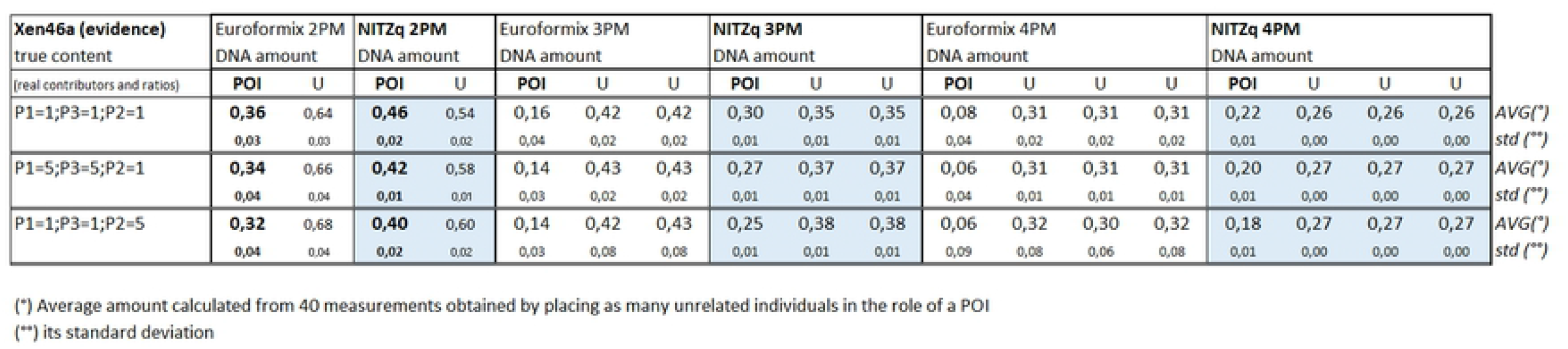
Average amount of DNA assigned to a cohort of 40 false contributors placed (one at the time) in the role of POI at a three-person-based experimental evidence (XEN46a)

**Fig 7.**
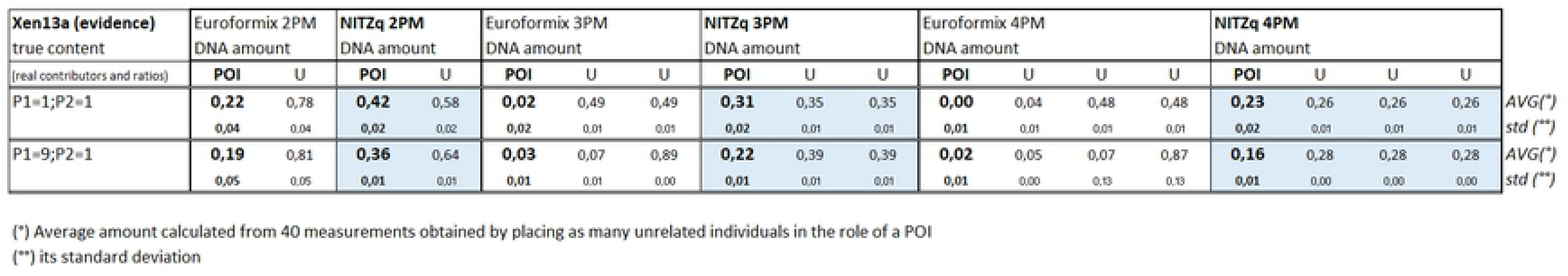
Amount of DNA assigned to a cohort of 40 false contributors placed (one at the time) in the role of POI at a two-person-based experimental evidence (XEN13a)

### LR values from true contributors as POI

Log10(LR) values calculated from the two tutorial datasets are in Fig 8 and Fig 9. To produce this data, we set NITZq to call the biallelic evidence every time the minor allele read would be ≥ to the major read multiplied by 0,05 and call the mono-allelic below this value. EUROFORMIX calculations were based on an absolute minor allele calling threshold of 5 reads and a P(D)= 0.05. Semi-continuous LRs are of comparable value if the two systems of calculation end up to call the same homozygous genotypes within the whole locus evidence. Betting on the wrong number of contributors does not significantly affect the semi-continuous LRs of either system. Both NITZq and EUROFOMIX return comfortably high values of quantitative LRs at tests where POIs match imbalanced evidence. However, NITZ underestimates LRs by 1 to sometimes 10 orders of magnitude (or bans), compared to EUROFORMIX, when the POI predominates in quantity within the evidence. To investigate about where such a difference can come from, we compared individual locus specific LRs obtained from selected mixture schemes to the corresponding single –source LR values (1/P(GT), or 1 divided by the POI’s genotype frequency). We consequently noticed that EUROFORMIX ver. 3.3. often exceeds the 1/P(GT) value, whereas NITZq never behaves like this. To cite an example, at a typical P1UU/UUU calculation based on XEN46a tutorial dataset ratio 5:1:1 (with P1= POI and P1 DNA quantity exceeding five times P3 and P2 in the mixture content) this happened at more than fifty over 133 overall loci (data not shown). According to Cowell et al [ 27] the weight of evidence (WOE) to draw from a suspect’s profile compared to a matching mixture cannot be stronger than that obtained when the suspect matches the single-source stain, or:

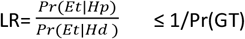

(where Pr(GT) is the suspect’s genotype frequency. We believe that this scores a strong point on NITZ’s side and earns credibility to its predictions.

**Fig 8.**
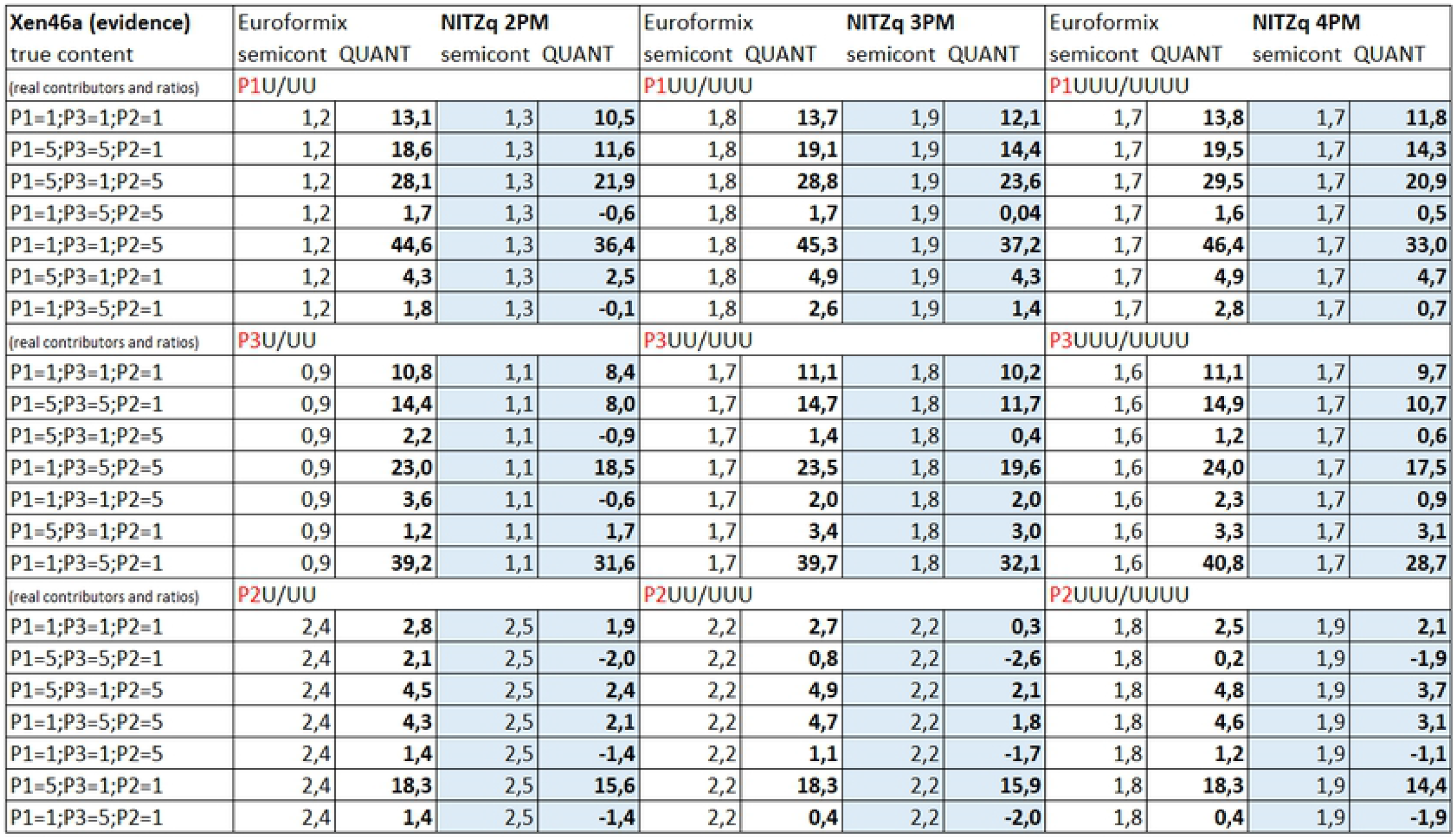
Log10(LRs) pertaining to three true contributors matched (one by one) as POI to an experimental three person mixture (XEN46a). EUROFORMIX and NITZq calculations are shown in comparison

**Fig 9.**
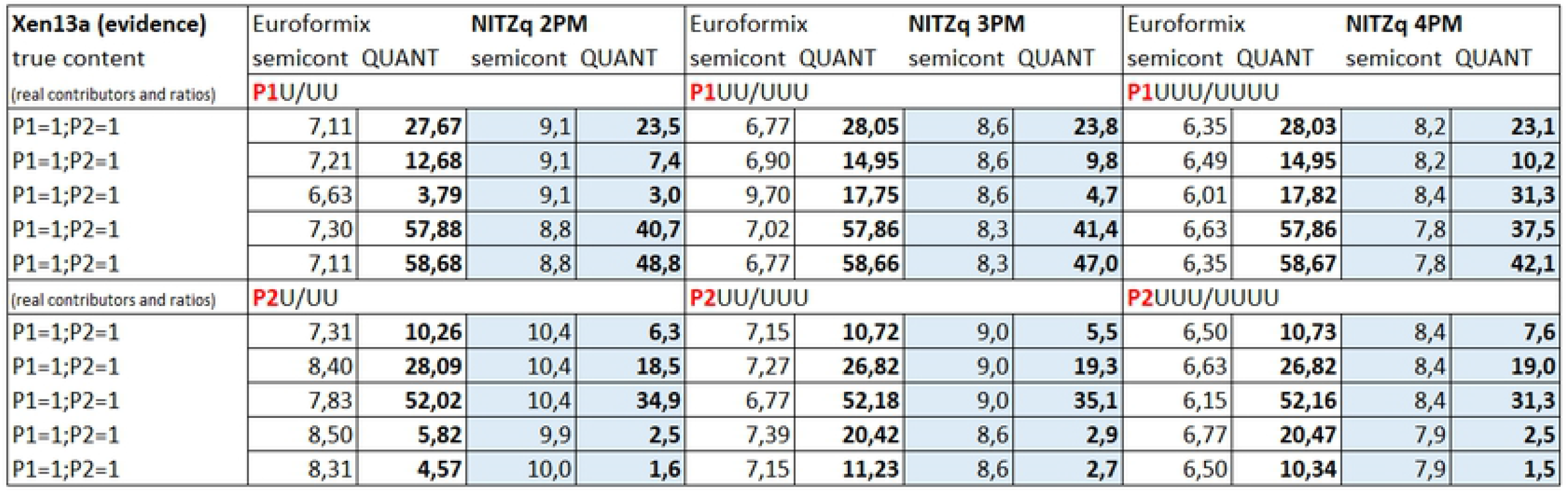
Log10(LRs) pertaining to three true contributors matched (one by one) as POI to an experimental two person mixture (XEN13a). EUROFORMIX and NITZq calculations are shown in comparison

### LR values for false contributors as POI

These datasets were produced under the same scheme of analysis detailed in the previous chapter. An excerpt of the whole data showing the maximum and minimum values of LRs, along with the internal percentile distribution is in Fig 10 and Fig 11, where NITZ and EUROFORMIX are shown in direct comparison (log10(LR) are shown). It is easily noticed that NITZq returns much lower LRs on false contributors, thus ruling out type-two errors (false inclusions) more efficiency than EUROFORMIX. This is essentially on account of three issues:

a. NITZ is highly efficient in spotting the homozygous evidence, thanks to the dynamic threshold (based on the minor vs major read ratio) used to call the minor read allele;
b. As a consequence, NITZ allows for more exclusion cases to emerge - with evidence E{a} either facing a POI=AB or a POI= BB;
c. NITZ deals with these two classes of exclusion by algebra, according to equ. 6-9; this results in low ((E{a} vs K=AB) or very low (E{a} vs K=BB) LR values.

**Fig 10.**
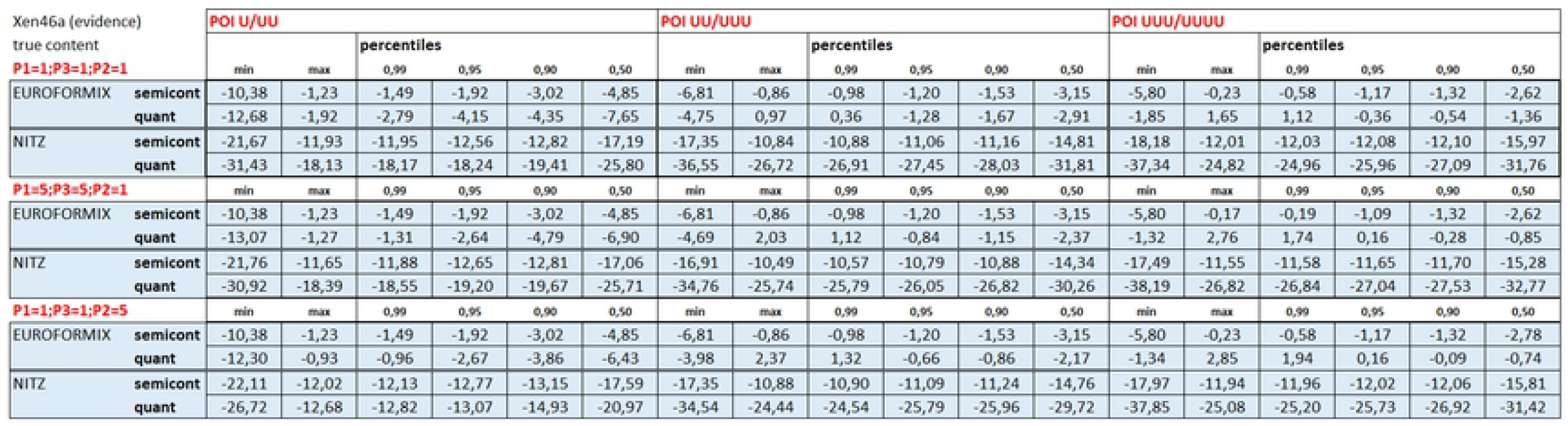
Log10(LRs) pertaining to forty false contributors matched (one by one) as POI to an experimental three person mixture (XEN46a). EUROFORMIX and NITZq calculations are shown in comparison

**Fig 11.**
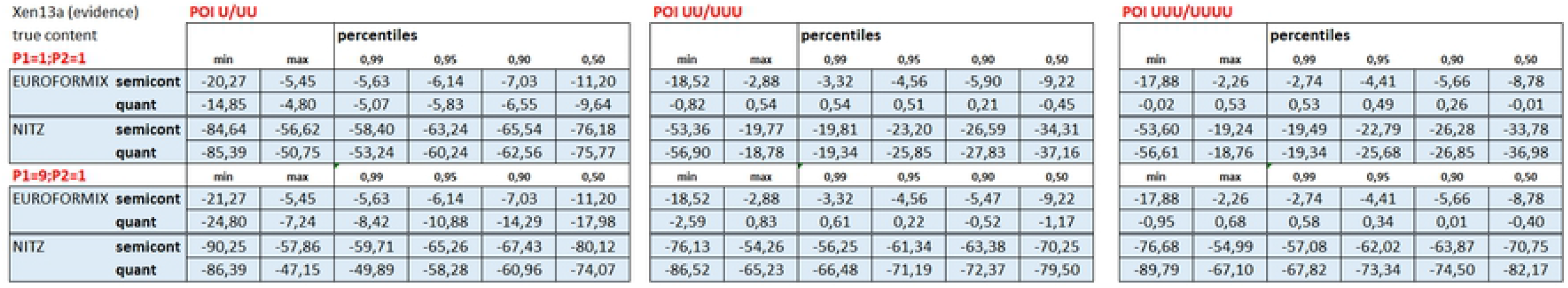
Log10(LRs) pertaining to forty false contributors matched (one by one) as POI to an experimental two person mixture (XEN13a). EUROFORMIX and NITZq calculations are shown in comparison

### Correlation between MR and LR values, with true contributors as POI

The LR/MR relationship was investigated at both the XEN46a and XEN13a evidence, by matching P1, P2 P3 (the ‘true’ contributors) as POI to variously imbalanced mixtures. For simplicity, LR values were calculated by adopting the true number of contributors (at XEN13a: KU/UU; at XEN46a: KUU/UUU). Then the Log10(LR_quant_) minus the Log10(LR_semicont_) value was paired to the quantitative fraction predicted to each POI. Pairs of values were plotted together, regardless of the difference in genetic profile originating the data-points.

We found that when true contributors are chosen as POI, a direct proportional relationship between the POI MR proportion and the LOG10(LR) value emerges from the NITZ-generated series of data (Fig 12; Fig 13). We concluded that the higher MR fraction of the POI the higher LR value. This relationship holds well even when a wrong number of contributors is used to calculate the LRs (data not shown). Data generated by EUROFORMIX behaved in the same way (data not shown).

**Fig 12.**
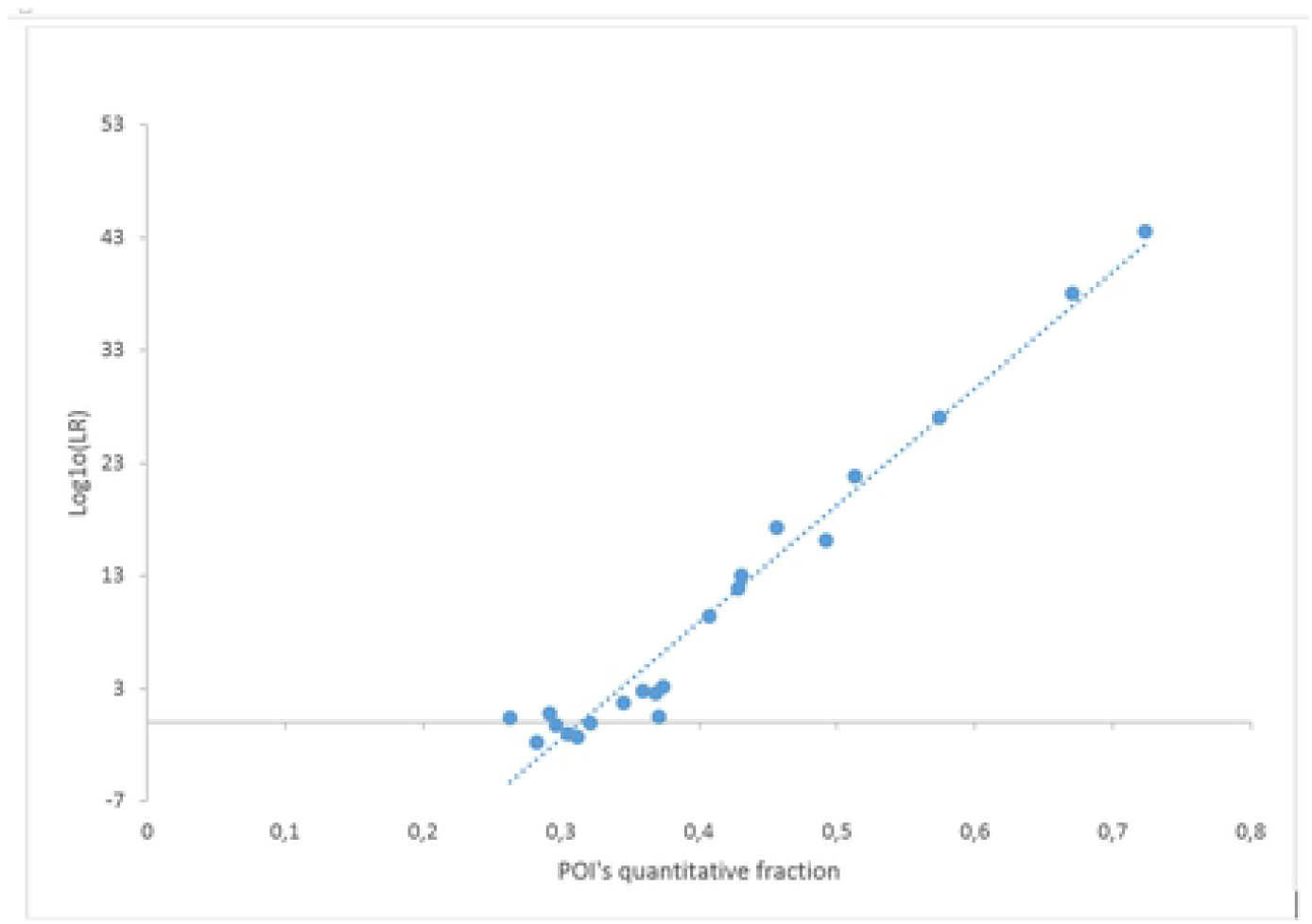
Plots of POI’s DNA amount vs Log10(LR) value in a series of true contributors tests. The evidence is XEN46_a, POIs are P1,P2,P3 (all downloaded from EUROFORMIX website). The NITZq 3PM worksheet was used to produce these results.

**Fig 13.**
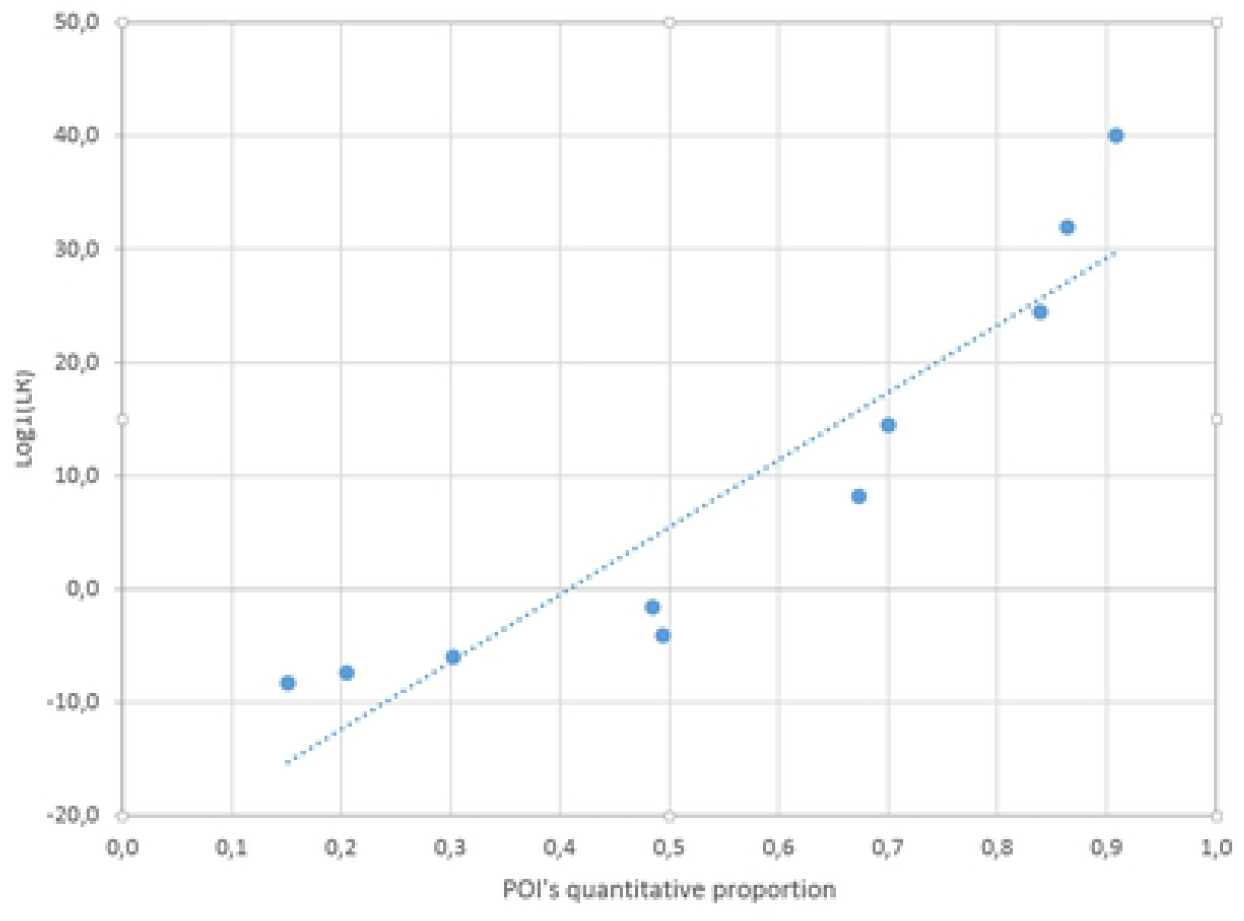
Plots of POI’s DNA amount vs Log10(LR) value in a series of true contributors tests. The evidence is XEN13_a, POIs are P1,P2 (all downloaded from EUROFORMIX website). The NITZq 2PM worksheet was used to produce these results.

### MR and LR relationship – with false contributors as POI

This relationship was investigated by the same scheme of analysis as adopted in the previous chapter. Forty false contributors were chosen – one at the time - as the POI in the following schemes of analyses: XEN46a ratio 1.1:1, KUU/UUU; XEN13a ratio 1.1, KU/UU. At false contributors tests, no coherent relationship was found among MR and le LR (Fig 14; Fig 15).

**Fig 14.**
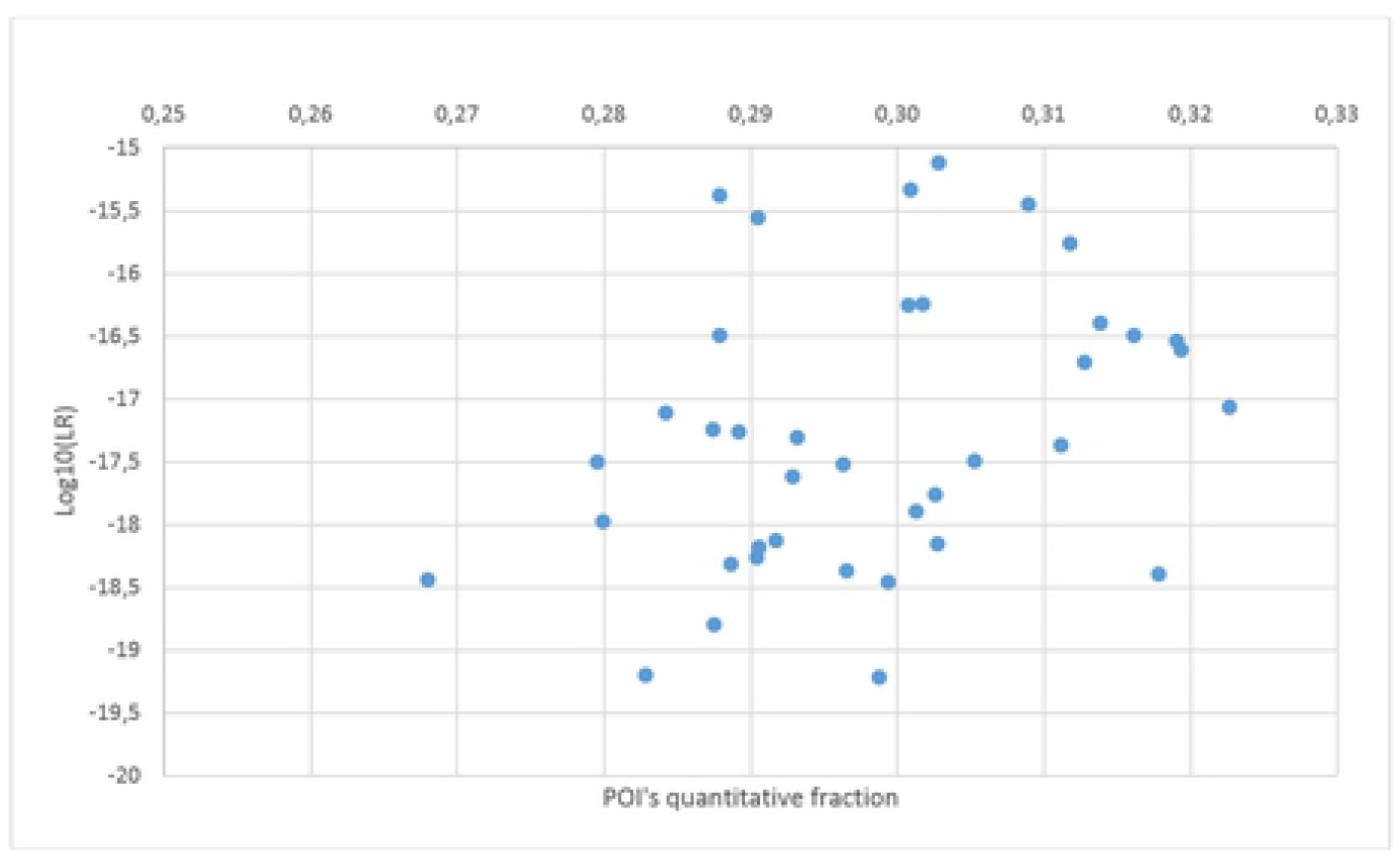
Plots of POI’s DNA amount vs Log10(LR) value in a series of false contributors tests. The evidence is XEN46_a ratio 1:1:1, POIs are forty unrelated individuals. The NITZq 3PM worksheet was used to produce these results.

**Fig 15.**
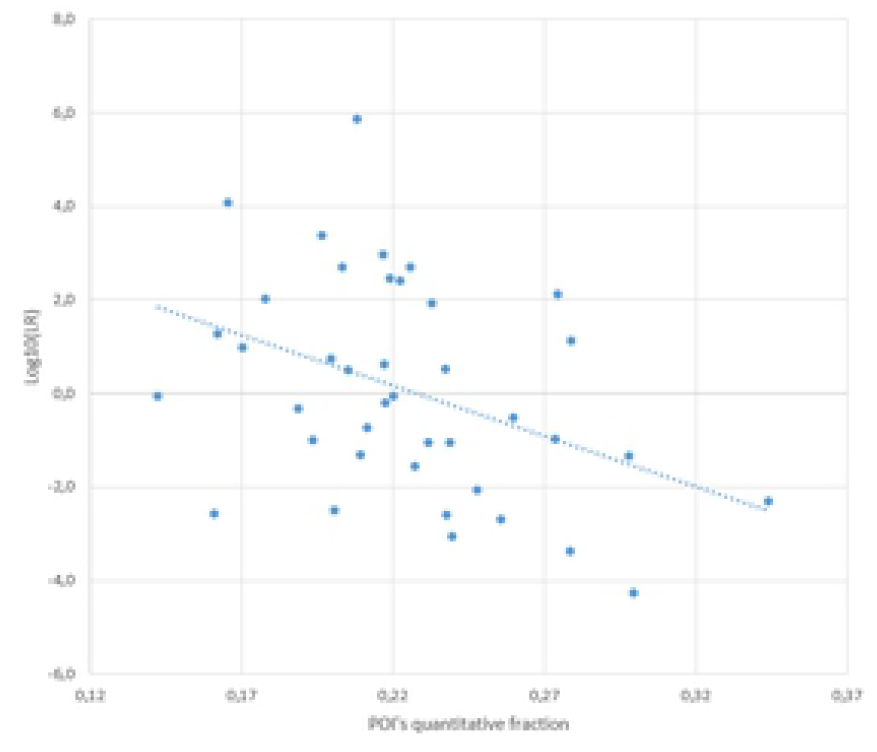
Plots of POI’s DNA amount vs Log10(LR) value in a series of false contributors tests. The evidence is XEN13_a ratio 1:1, POIs are forty unrelated individuals. The NITZq 2PM worksheet was used to produce these results.

## Discussion

We have introduced a series of original procedures meant to re-define the way semi-continuous and quantitative calculations are carried out at binomial markers.

On the **semi-continuous** side of the issue, we vindicate the viewpoint that the routine binomial evidence at a mixture cannot be but bi-allelic (Et{a;b}) and that this predominant example of SNPs evidence is quite well modelled by matrices of multiple, equally possible genotypic permutations. Within these discrete framework of probability matrices, all genotypes entering in the same permutation state multiply together and all permutation states products sum up, so as to give the true value of a likelihood. As long as calculations rest on only genotype probabilities, the matrix scheme of calculation can be easily reduced to algebra - and we have here introduced a series of notations that make previously reported formulas [34] of universal use in the semi-continuous context. Interestingly, matrix-based calculations and algebraic calculations give identical results, thus implying that the two underlying methods (matrices; algebra) are sound and each is consistent to the other. If binomial matrices and algebraic formulas are used, semi-continuous inference at SNPs is reduced to the application of simple arithmetic and it does not require precompiled software. This way to calculate does not undergo any limitation in the number of contributors and in the number of POIs to compute on- as often dedicated software does.

In turn, we interpret mono-allelic evidence at a SNPs mixture (Et{a}) as a ‘residual’ evidence – the product of a biallelic evidence having undergone dropout events. We assign the Et{a} locus evidence only when the h index at a given locus falls below a predetermined, dynamic threshold reflecting extreme imbalance among the minor and the major read of an individual NGS record. Albeit residual in their practical occurrence, examples of the monoallelic evidence Et{a} emerging at a SNPs protocol can modify the inferential value of an identity test to a big extent, by either considerably increasing the LR (case: Et{a} /POI_AA_) or considerably lowering it (cases: Et{a} /POI_BB_; Et{a} /POI _BB_). To calculate the relevant LRs at these ‘residual but important’ cases we have worked out nine algebraic solutions covering all possible instances of Et{a}/POI genotype assortment. Within these formulas the ‘h’ index – or the minor vs major read ratio of a typical NGS experiment – plays the role of an acceptable proxy of the dropout probability, and no experimental P(DO) values are needed. The whole procedure just here summarized becomes, in our view, an acceptable equivalent of using the split-drop approach.

As far as the **quantitative** calculation is concerned, we here introduce an empirical way to capture the ratio at which any give POI of interest mix with the other unknown mixture contributors. We examine all permutations containing the POI- genotype at all loci of any standard testing protocol of choice and, among these, we discern/extract all those that do not involve allele overlaps between the POI’s genotype and the unknowns’ (NOCs) - along with all further cases involving very simple POI-Unknowns overlaps (SOCs). By simple arithmetic we extract the empirical POI share at all of these privileged loci, we average all values and we determine an expected ‘POI-vs-Unknowns’ ratio. Compliance between this unique expected values and all observed MR values pertaining to all existing deconvolution states (measured as a (1-χ^2^) value) gives us the probability of any permutations states to exist on the quantitative side.

Compared to other methods [23–55] our method for calculating the mixture MR is surprisingly simple. Its computation within a spreadsheet is practically instantaneous and it ends up to give back quantitative ratios that approximate well the MCMC and MLE estimates.

Unlike other methods, our MRs flatten the Unknwns share to the same figure, due to the phenomenon of quantity permutations across the matrix. NOCs-SOCs based (1-χ^2^) values are true probability values and they are independent of the genotype probabilities and the h indexes. These three independent probabilities are therefore multiplied together to yield the value of a likelihood, according to a unique procedure of our conception ( ‘NITZq’ procedure).

We extensively tested NITZq by choosing a 133 c SNPs loci protocol [actually, a subset of the 140 loci panel of Grandell eta l 2016], by using the “dataset 4: SNPs data” package available at the EUROFORMIX website as evidence and by assuming the relevant EUROFORMIX calculations as the benchmark. We here show that when true contributors are tested as POI, NITZq computations return LR values of comparable magnitude as those returned by EUROFORMIX. Type-one errors (mediocre or negative LOGLRs) occasionally occur with NITZ – essentially when the true contributor’s quantitative share tends to disappear from the mixture Additionally, when the POI is a true contributor and is quantitatively predominant, NITZ usually gives back a lower value of the evidence (WOE) than EUROFORMIX. As an explanation of this latter phenomenon, we report that the EUROFORMIX individual LRs often exceed the corresponding single-source WOE (namely: one over the relevant genotype frequency value) – usually regarded as a ceiling value not to be crossed while evaluating the presence of individual profiles within a mixture. NITZq never oversteps the relevant single-source WOE and it therefore qualifies as a fully realistic approach to the quantitative analysis. When false contributors are chosen as POI, a much lower LR is gathered by NITZq, compared to EUROFORMIX. Type-two errors (false inclusions) occasionally occur with EUROFORMIX whereas the practically do not occur with NITZq. We believe that this is due to the circumstance that NITZ spots incompatible events much more efficiently than EUROFORMIX. When a true contributor is a POI, a linear relationship exists among its quantitative share in the mixture and the log10(LR) value, as calculated by NITZ. When the POI is a false contributors, this relationship becomes incoherent and graphically sparse. With Nitz, betting on the wrong number of contributors does not necessarily result in lower LRs. So the rule of three invoked by Bleka et al [36] has limited application with NITZ.

## Data Availability Statement

The MS Excel files cited in this article are freely available at:

www.vincepascali.it/docs/NITZsc_2_to_8_contrs.xlsx
www.vincepascali.it/docs/NITZq_SNPs_2PM.xlsx
www.vincepascali.it/docs/NITZq_SNPs_3PM.xlsx
www.vincepascali.it/docs/NITZq_SNPs_4PM.xlsx

